# Unlocking the *Alphitobius diaperinus* (Coleoptera: Tenebrionidae)-combatting capabilities of *Bacillus thuringiensis* INTA Mo4-4 through genomic and phenotypic characterization

**DOI:** 10.64898/2026.05.13.724709

**Authors:** Melisa Paula Pérez, Leopoldo Palma, Marcelo Facundo Berretta, Graciela Beatriz Benintende, Diego Herman Sauka

## Abstract

*Bacillus thuringiensis* INTA Mo4-4 was characterized phenotypically, genomically, and for insecticidal activity against *Alphitobius diaperinus*. Microscopy revealed rare flat rectangular parasporal crystals, and SDS-PAGE identified a *ca*. 67 kDa protein, similar to *B. thuringiensis* serovar *morrisoni* strain *tenebrionis* DSM-2803, which was proteolytically processed to a *ca*. 55 kDa fragment. Genomic analysis showed a 5.99 Mb genome with 99.43% completeness, clustering phylogenetically with *B. cereus* and *B. thuringiensis*. High genomic similarity was observed with *B. thuringiensis* svar. *morrisoni* BGSC 4AA1, confirmed by MLST analysis assigning it to ST-23. The genome encodes an interesting arsenal of pesticidal proteins showing significant similarity to Cry3Aa, Mpp23Aa, Xpp37Aa, Mpp5Ab, Vpb1Ad, Vpb1Ae, Vpa2Ab, Vpa2Ba, Vpa2Bb and Spp1Aa, with demonstrated toxicity against coleopteran pests. Biosynthetic gene clusters for toyoncin, fengycin, and bacillibactin were identified. Dose-response bioassays showed that INTA Mo4-4 was nearly four times more toxic to A. diaperinus larvae (LC_50_ 136.9 µg/ml) than DSM-2803 (LC_50_ 540.5 µg/ml), with the difference being statistically significant. No teratological effects were observed on *Musca domestica*. These findings suggest that INTA Mo4-4 is a promising candidate for the biological control of *A. diaperinus*.

## 1. Introduction

*Bacillus thuringiensis* is a Gram-positive, spore-forming bacterium that has become a cornerstone of biological pest control due to its remarkable efficacy against a wide range of insect pests from the orders Lepidoptera, Diptera, and Coleoptera. This versatility is attributed to its production of virulence factors, including crystalline (Cry and Cyt) proteins synthesized during sporulation and vegetative insecticidal proteins such as Vip3 and binary Vpa and Vpb proteins, secreted during the vegetative growth phase [1]. These proteins specifically target insects upon solubilization in the alkaline midgut, where they are proteolytically activated by gut enzymes before binding to receptors on the brush border membrane of epithelial cells, leading to pore formation, cell lysis, and insect death [2]. Additionally, some *B. thuringiensis* strains can secrete thermostable non-proteinaceous toxins, such as thuringiensin (also known as Type I β-exotoxin), during vegetative growth [3]. However, unlike Cry proteins, thuringiensin lacks high specificity, raising concerns about its potential toxicity to non-target organisms and limiting its suitability for biocontrol applications [3].

The specificity and efficacy of different *B. thuringiensis* strains can vary significantly, driving ongoing research to identify and characterize novel strains with enhanced toxicity profiles and broader host ranges [4,5]. Advances in genomic analyses have greatly contributed to this effort by identifying genes responsible for virulence and host specificity, facilitating the discovery of more effective strains and the development of targeted bioinsecticides [3,6].

Among agricultural pests of significant concern, *Alphitobius diaperinus* (Coleoptera: Tenebrionidae), commonly known as the lesser mealworm, poses a major threat to poultry production systems. This pest acts as a vector for pathogens that compromise poultry health, causing traumatic wounds such as feather and skin lesions in chicks, increasing their susceptibility to infections, and accelerating the spread of diseases, including bacteria and parasites, among chicken stocks [7]. These effects lead to substantial economic losses for poultry producers. Additionally, the beetle poses occupational risks to farm workers through exposure to contaminated materials and beetle-related allergens, adding to the challenges in poultry production [7].

Traditional control strategies for *A. diaperinus* rely on chemical insecticides applied during sanitary downtime between production cycles to prevent harmful residue exposure. However, insecticide resistance and increasing environmental concerns have highlighted the need for alternative control approaches [8]. In this context, *B. thuringiensis*-based biopesticides represent a promising solution to control *A. diaperinus* throughout the entire poultry production process due to their high specificity, low environmental impact, and demonstrated safety for non-target organisms, including poultry [9,10,11].

As part of a screening program of *B. thuringiensis* strains native to Argentina, the INTA Mo4-4 strain was selected for its high toxicity against *A. diaperinus* larvae. Among the forty-one strains evaluated, it exhibited the highest insecticidal activity [12]. Isolated from the Bacterial Collection of the Instituto de Microbiología y Zoología Agrícola (IMYZA-INTA, Argentina), this strain caused mortality rates 2.7 times higher than those produced by the well-known *B. thuringiensis* serovar *morrisoni tenebrionis* DSM2803 in preliminary bioassays, a reference strain effective against various coleopteran pests [12]. Although some evidence suggests that INTA Mo4-4 may produce insecticidal proteins during vegetative growth, its primary virulence factors are crystal proteins [12].

This study aims to perform a comprehensive genomic and toxicological characterization of the INTA Mo4-4 strain to elucidate its insecticidal mechanisms against *A. diaperinus*, supporting its potential development as a novel *B. thuringiensis*-based biopesticide for coleopteran pest management.

## 2. Results

### 2.1. Strain characterization

Observations of the spore-crystal complex under phase-contrast microscopy revealed that INTA Mo4-4 exclusively produces flat, rectangular parasporal crystals within mature sporangia (Figure 1A). In addition, scanning electron microscopy (SEM) confirmed the release of both spores and crystals and showed that the crystals exhibit size variability, with average dimensions of 1.329 × 0.985 μm (Figure 1B).

**Figure 1.**
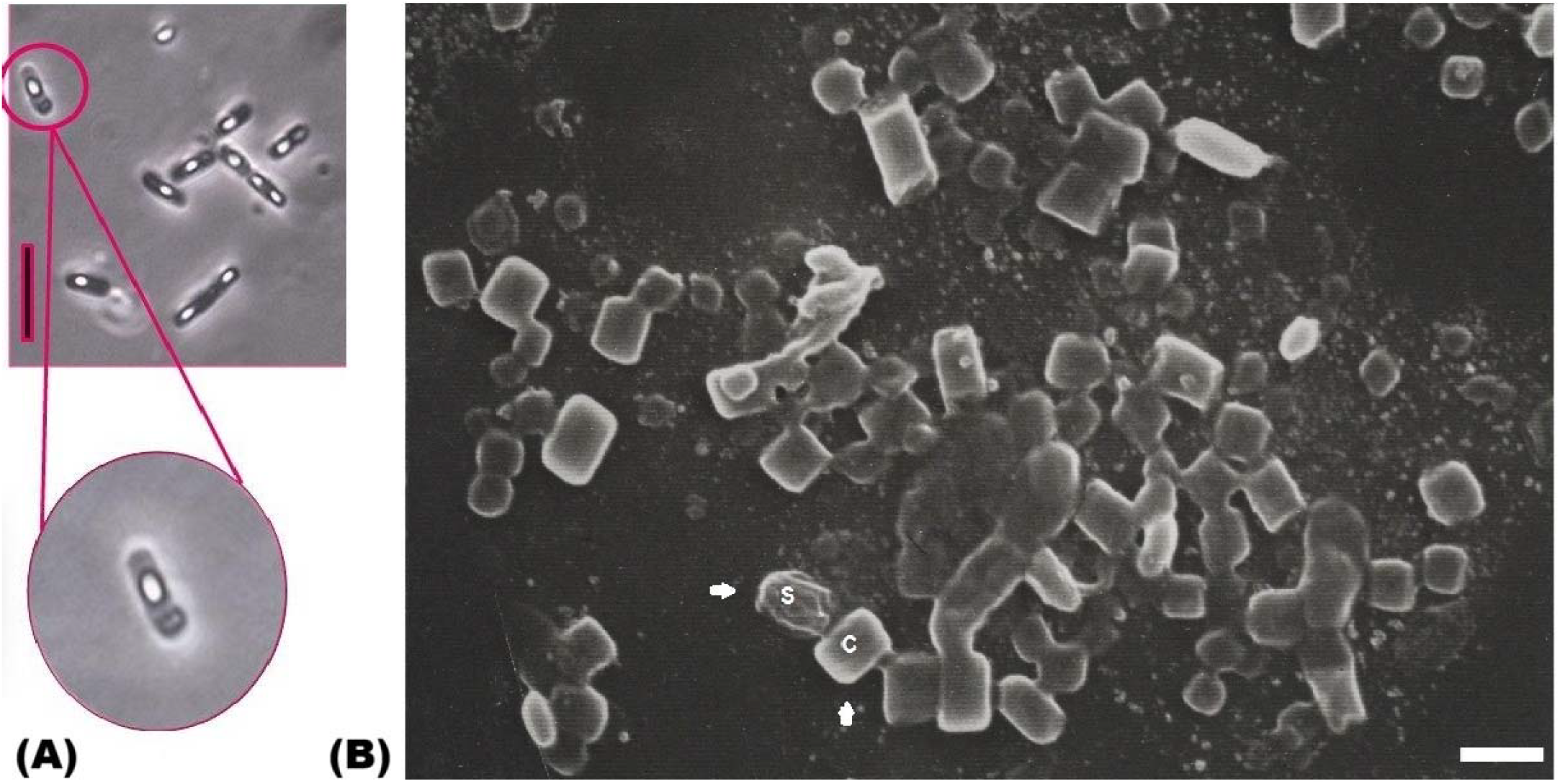
Microscopic analysis of INTA Mo4-4. Flat rectangular parasporal crystals are observed within mature sporangia by phase-contrast microscopy (A, scale bar: 5 μm), and released crystals (C) and spores (S) are shown by SEM (B, scale bar: 1 μm).

SDS-PAGE analysis of the spore-crystal complex revealed a prominent band with an approximate molecular weight of 67 kDa (Figure 2A). This band co-migrated with the major protein band from *B. thuringiensis* svar. *morrisoni* strain *tenebrionis* DSM-2803. Additionally, to assess the solubility and proteolytic activation of the crystals, the spore-crystal complex of INTA Mo4-4 was tested under different pH conditions (Fig. 2B) and subjected to trypsin digestion. SDS-PAGE analysis of the trypsin-digested samples revealed co-migrating protease-resistant peptides of *ca*. 55 kDa (Figure 2C).

**Figure 2.**
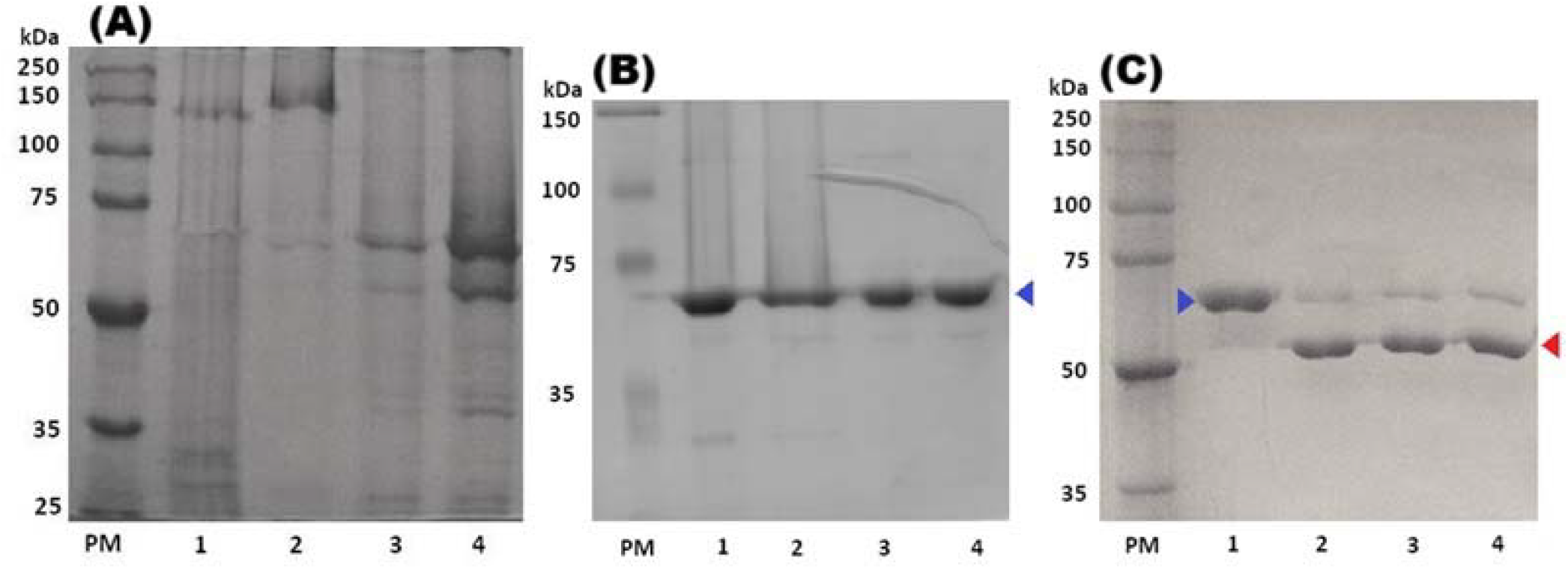
(A) Electrophoretic analysis of proteins forming the crystals. Lanes: 1, *B. thuringiensis* svar. *israelensis* IPS-82; 2, *B. thuringiensis* svar. *kurstaki* HD-1; 3, *B. thuringiensis* svar. *morrisoni tenebrionis* DSM2803; 4, INTA Mo4-4. (B) Electrophoretic analysis of solubilized crystal proteins and (C) solubilized proteins digested with trypsin from INTA Mo4-4. (B) Lanes: 1, no treatment; 2, NaBr 3.3 M, pH 6.6; 3, NaOH 0.1 M, pH 12.9; 4, Na_2_CO_3_ 50 mM, pH 10.6. (C) Lanes: 1, solubilized crystal proteins (Na_2_CO_3_ 50 mM, pH 10.6); 2, 2 h incubation; 3, 4 h incubation; 4, 6 h incubation. The blue arrows indicate bands corresponding to the solubilized proteins (*ca*. 67 kDa), while the red arrow highlights the solubilized and trypsinized proteins (*ca*. 55 kDa). Protein markers (PM) with sizes indicated on the left (kDa) (Promega).

### 2.2. Genomic characterization and comparative analysis

The draft genome assembly of the *Bacillus* sp. strain INTA Mo4-4 was evaluated for quality and completeness using QUAST v5.3.0 and CheckM v1.0.18. The assembly consisted of 101 contigs, totaling 5,988,741 nucleotides, with a GC content of 34.88%. It predicted 6,454 protein-coding sequences (CDS) and 66 RNA genes. The largest contig measured 363,546 nucleotides, and the assembly achieved an N_50_ of 134,295 and an L_50_ of 14. The genome exhibited a high completeness of 99.43%, with no detectable contamination.

Further analysis using BUSCO v5.8.0 confirmed functional completeness, identifying all 302 orthologues from the bacilli_odb10 database as complete (100%), with 99.3% as single-copy genes. A similar analysis with the bacillales_odb10 database identified 447 orthologues (99.8%) as complete, with 98.9% as single-copy genes.

Phylogenetic analysis using the Type Strain Genome Server (TYGS) placed INTA Mo4-4 in a distinct cluster, closely related to the type strains *B. cereus* ATCC 14579 and *B. thuringiensis* ATCC 10792, suggesting that INTA Mo4-4 may represent a novel species (Figure 3).

**Figure 3.**
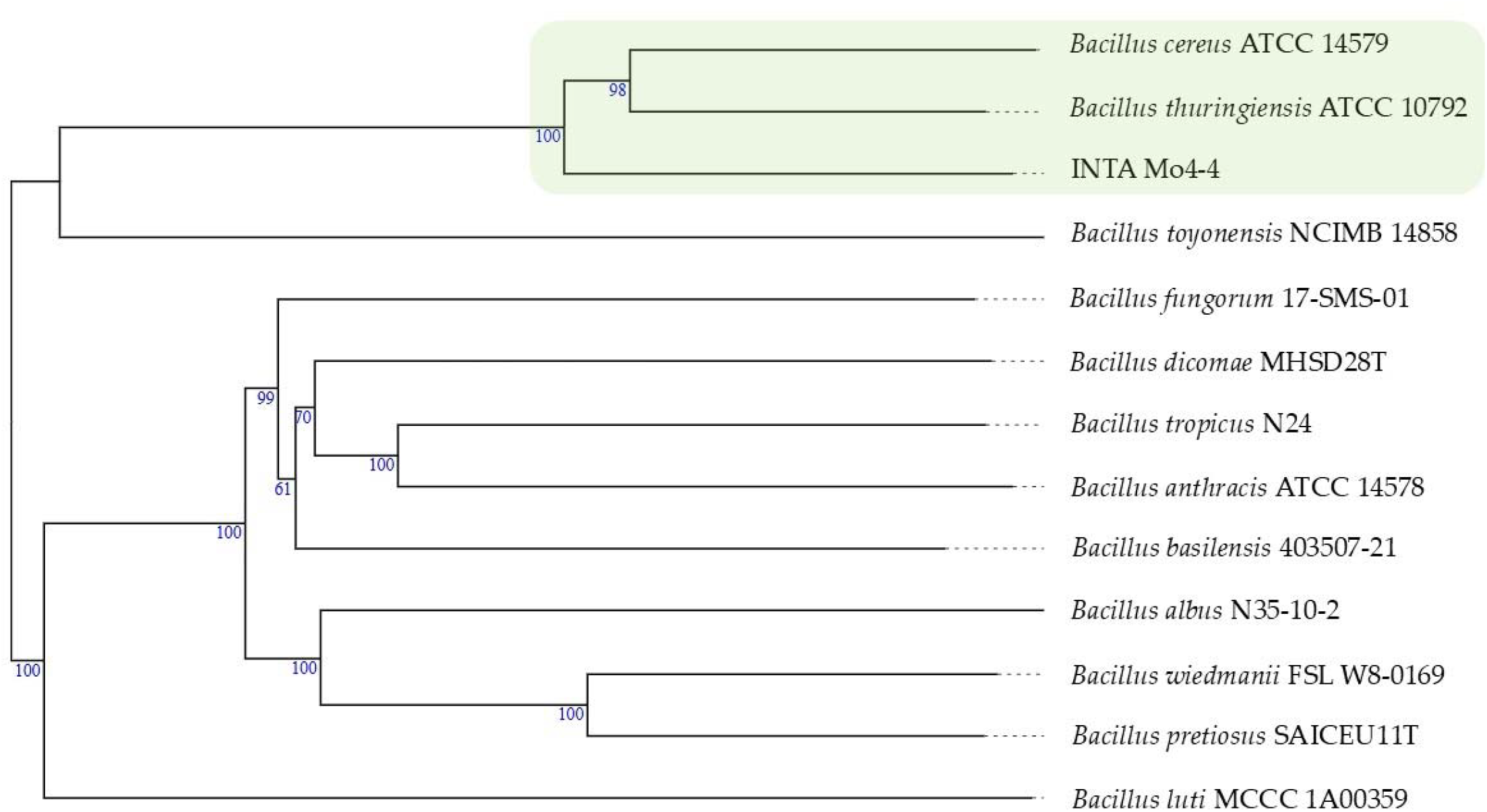
GBDP phylogenetic tree generated from whole-genome data using the TYGS, with an average branch support of 92.8%. The green color is used to highlight the clustering of INTA Mo4-4 with the type strains.

Genomic similarity at the nucleotide level between INTA Mo4-4 and the 12 most closely related strains is presented in Table 1. The comparison reveals a high degree of similarity, particularly with *B. thuringiensis* svar. *morrisoni* strain BGSC 4AA1, which exhibited the highest values across all metrics (Table 1). Other strains, including *B. thuringiensis* and *B. cereus* type strains, also showed significant similarity, with ANIb and ANIm values exceeding 95 % and consistently high tetra-nucleotide z-scores (Table 1).

**Table 1.**
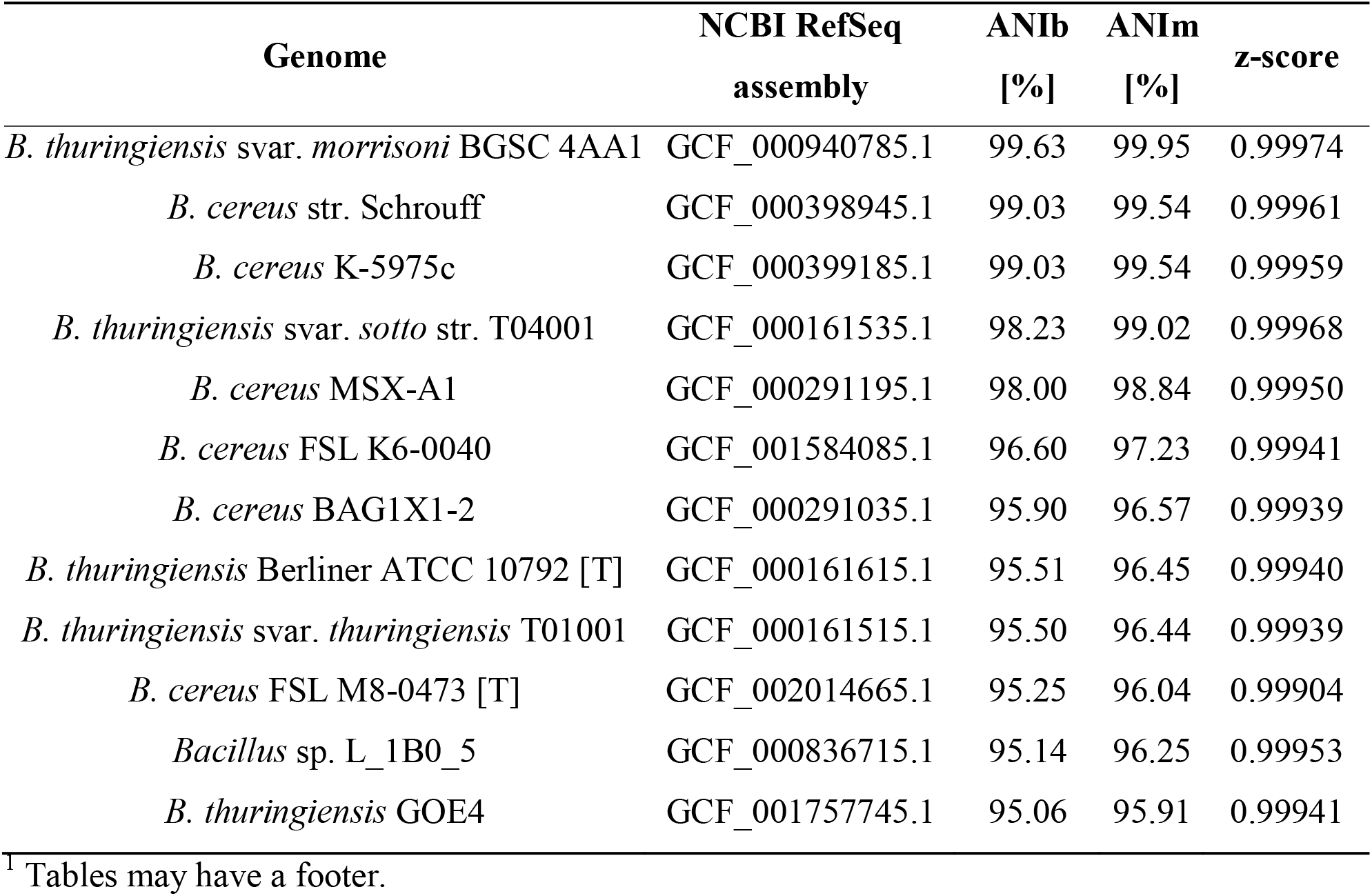
Genomic similarity at the nucleotide level between INTA Mo4-4 and other closely related strains. Metrics include average nucleotide identity by BLAST (ANIb), average nucleotide identity by MUMmer (ANIm), and tetra-nucleotide z-scores [13]. [T] Type strains.

| Genome | NCBI RefSeq<br>assembly | ANiB<br>[%] | ANIm<br>[%] | z-score |
| --- | --- | --- | --- | --- |
| *B. thuringiensis* svar. *morrisoni* BGSC 4AA1 | GCF_000940785.1 | 99.63 | 99.95 | 0.99974 |
| *B. cereus* str. Schrouff | GCF_000398945.1 | 99.03 | 99.54 | 0.99961 |
| *B. cereus* K-5975c | GCF_000399185.1 | 99.03 | 99.54 | 0.99959 |
| *B. thuringiensis* svar. *sotto* str. T04001 | GCF_000161535.1 | 98.23 | 99.02 | 0.99968 |
| *B. cereus* MSX-A1 | GCF_000291195.1 | 98.00 | 98.84 | 0.99950 |
| *B. cereus* FSL K6-0040 | GCF_001584085.1 | 96.60 | 97.23 | 0.99941 |
| *B. cereus* BAG1X1-2 | GCF_000291035.1 | 95.90 | 96.57 | 0.99939 |
| *B. thuringiensis* Berliner ATCC 10792 [T] | GCF_000161615.1 | 95.51 | 96.45 | 0.99940 |
| *B. thuringiensis* svar. *thuringiensis* T01001 | GCF_000161515.1 | 95.50 | 96.44 | 0.99939 |
| *B. cereus* FSL M8-0473 [T] | GCF_002014665.1 | 95.25 | 96.04 | 0.99904 |
| *Bacillus* sp. L_1B0_5 | GCF_000836715.1 | 95.14 | 96.25 | 0.99953 |
| *B. thuringiensis* GOE4 | GCF_001757745.1 | 95.06 | 95.91 | 0.99941 |

<sup>†</sup> Tables may have a footer.

In silico multilocus sequence typing (MLST) analysis revealed that the INTA Mo4-4 strain belongs to sequence type 23 (ST-23), the same as *B. thuringiensis* svar. *morrisoni* BGSC 4AA1, based on the allele profiles of seven housekeeping genes: *glp* (allele 15), *gmk* (allele 7), *ilv* (allele 7), *pta* (allele 2), *pur* (allele 5), *pyc* (allele 8), and *tpi* (allele 13).

A total of 6,516 new genomic features were identified in the assembled genome using RASTtk— v1.073, of which 174 were classified as non-coding elements. Among the annotated features, 6,342 were predicted as coding genes (CDS), while non-coding elements included 111 non-coding repeats and sixty-three non-coding RNAs (Supplementary Data S1).

The draft genome sequence of INTA Mo4-4 contains potential pesticidal proteins and virulence factors associated with insect pathogenesis. Ten CDS showed significant BlastX similarity to known pesticidal proteins such as Cry3Aa, Mpp23Aa, Mpp5Ab, Xpp37Aa, Spp1Aa, Vpb1A, Vpa2A and Vpa2B which demonstrated efficacy against a broad spectrum of insects spanning five different orders (Table 2). Additionally, INTA Mo4-4 harbors other coding sequences encoding four putative chitinases and four chitin-binding proteins. Genes associated with thuringiensin biosynthesis, such as *thuE*, were absent.

**Table 2.**
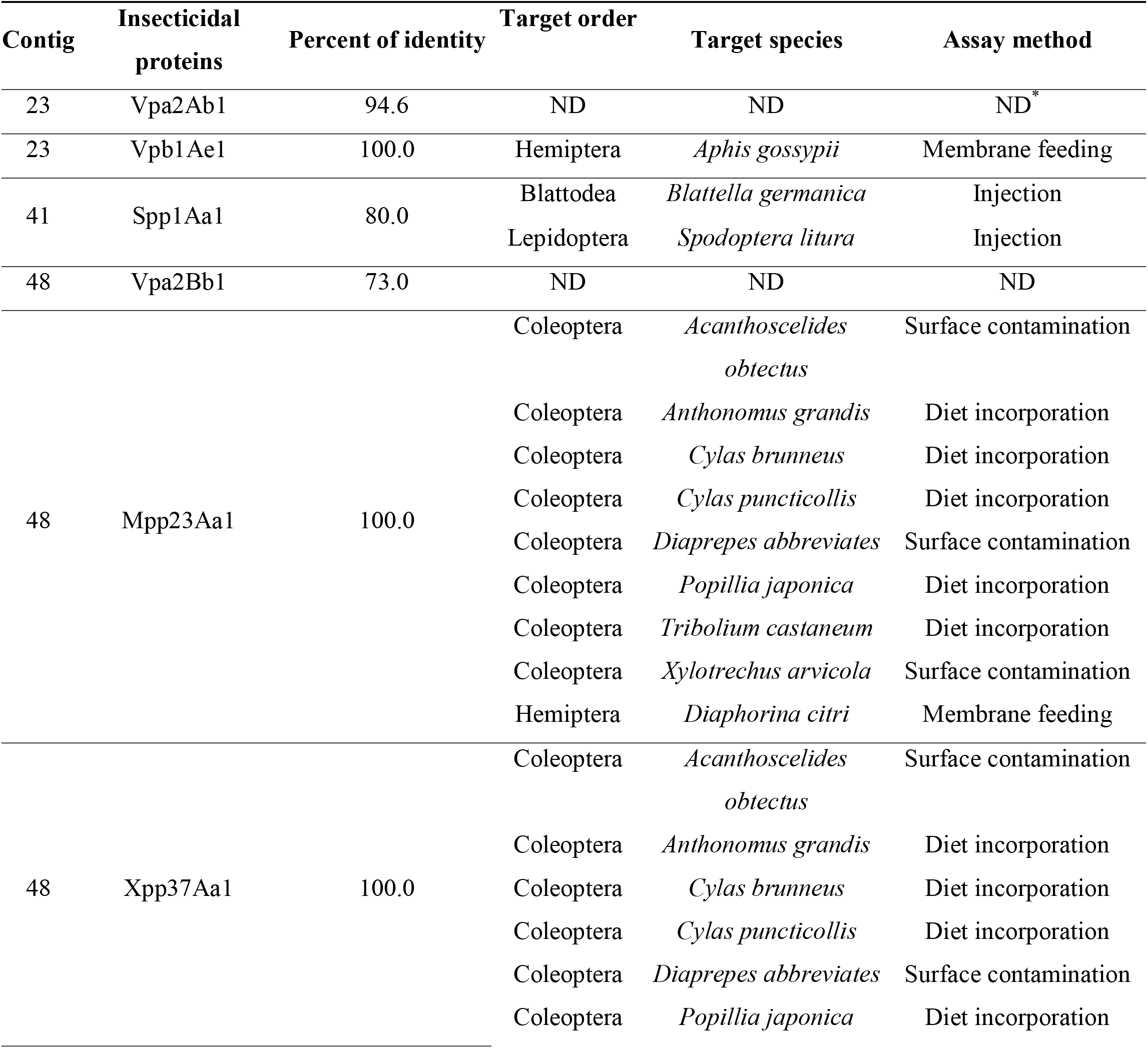
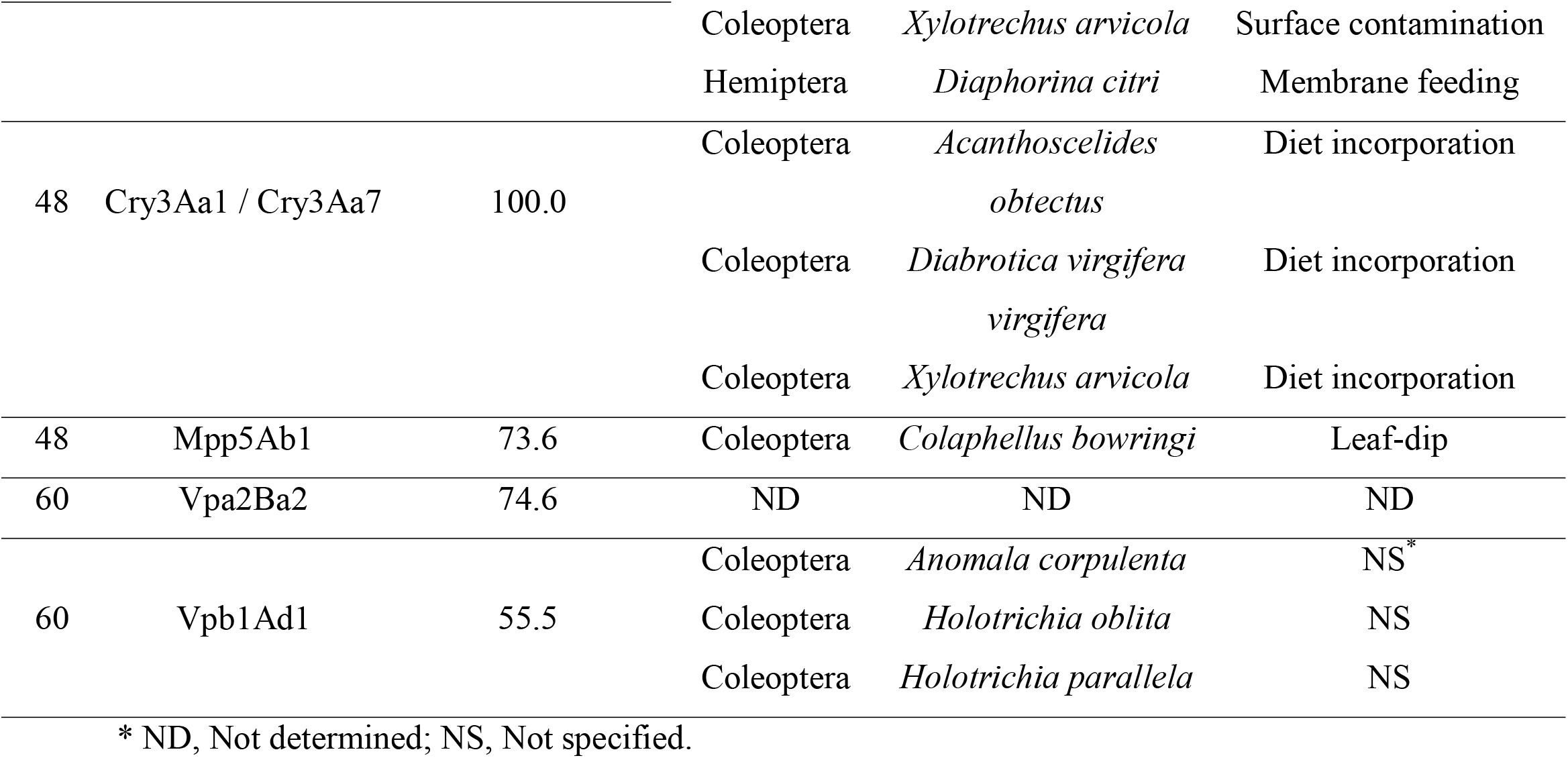
List of pesticidal proteins identified in the genome of INTA Mo4-4, using the specificity database containing experimental data from the Bacterial Pesticidal Protein Resource Center (BPPRC) for target species [14].

The genome of INTA Mo4-4 was further analyzed using the antiSMASH v6.1.1 tool, revealing several biosynthetic gene clusters (BGCs) with potential insecticidal and antimicrobial properties. A toyoncin-producing cluster, a ribosomally synthesized and post-translationally modified peptide (RiPP), was identified with 18% similarity to known references. Other BGCs include the synthesis of fengycin (40% similarity), a lipopeptide with antifungal activity, and bacillibactin (85% similarity), a siderophore with antimicrobial properties. A cluster with 100% similarity to petrobactin and various non-ribosomal peptide synthetase (NRPS) clusters were also detected. A complete list of the identified BGCs is provided in supplementary data S2.

### 2.3. Bioassays

Concentration-dependent bioassays were conducted to estimate the mean lethal concentrations (LC_50_) of INTA Mo4-4 and the reference strain *B. thuringiensis* svar. *tenebrionis* DSM2803 using Probit analysis. The results showed that the spore-crystal suspension of INTA Mo4-4 exhibited significantly higher toxicity than the reference strain, as evidenced by the non-overlapping confidence intervals (Table 3). Specifically, INTA Mo4-4 was 3.95 times more toxic than the *tenebrionis* DSM2803 strain.

Probit analysis was also performed on the active biomass (pellet) of INTA Mo4-4 subjected to two additional treatments: solubilization with Na_2_CO_3_ (50 mM, pH 10.6) and solubilization followed by trypsinization. The LC_50_ values (with 95% confidence limits) for these treatments were statistically indistinguishable from those of the untreated spore-crystal suspension, as indicated by overlapping confidence intervals (Table 3).

**Table 3.** Concentration-response insecticidal activity of spore-crystal suspensions of *B. thuringiensis* strains against second-instar larvae of *A. diaperinus* under untreated and treated conditions.

| Treatment/Strain | LC<sub>50</sub> (µg/ml)<sup>1</sup> | CV (%)<sup>2</sup> | Slope | χ<sup>2</sup> (4 df)<sup>3</sup> |
| --- | --- | --- | --- | --- |
| Untreated *tenebrionis* DSM2803 | 540.5 (334.3-1838.9) | 13.5 | 1.4 | 4.0 |
| Untreated INTA Mo4-4 | 136.9 (113.3-171.2) | 17.3 | 0.4 | 4.1 |
| Solubilized pellet INTA Mo4-4 | 158.4 (117.7-251.0) | 16.3 | 1.6 | 3.2 |
| Solubilized and trypsinized pellet INTA Mo4-4 | 164.5 (125.1-246.2) | 12.9 | 1.7 | 4.4 |

<sup>1</sup> Mean lethal concentration. Numbers in parenthesis are 95% confidence limits. <sup>2</sup> Coefficient of variation. <sup>3</sup> Chi-square.

Additionally, no teratological effects were observed during the emergence of *Musca domestica* Linnaeus (Diptera: Muscidae) adults from pupae.

## 3. Discussion

Research on *B. thuringiensis* interactions with coleopteran species is limited compared to other insect orders (*e.g*., Lepidoptera), which highlights the need for further investigation in this area. This gap emphasizes the potential for discovering new *B. thuringiensis* strains or proteins targeting coleopteran pests, which may offer new opportunities for biopesticide development [15].

This study provides a comprehensive genomic and phenotypic characterization of INTA Mo4-4, a promising biocontrol strain targeting *A. diaperinus*. Our results highlight the genetic similarity of this strain to *B. thuringiensis* svar. *morrisoni* BGSC 4AA1 and its enhanced insecticidal activity against *A. diaperinus*, suggesting its potential as an active alternative to conventional chemical insecticides. In addition, our findings elucidate key mechanisms underlying its pesticidal activity, highlighting the potential of INTA Mo4-4 as a candidate for biocontrol applications.

The morphological analyses of INTA Mo4-4 revealed the presence of rare, flat rectangular parasporal crystals. This morphology aligns with the crystalline features observed in *B. thuringiensis* svar. *morrisoni* DSM-2803, which is known for its specificity against coleopteran pests [16]. The identification of a prominent *ca*. 67 kDa protein band, which proteolytically degrades to a stable *ca*. 55 kDa form, along with the encoded *cry3Aa1* gene, strongly suggests the presence of expressed Cry3-like proteins in the crystal inclusions. Such proteins are well-documented for their insecticidal activity against coleopterans [14], supporting the hypothesis that INTA Mo4-4 operates through a similar mode of action.

The ability of the crystals to solubilize under alkaline conditions (pH 10.6) and the proteolytic stability of the resulting peptides are critical for their efficacy within the alkaline midgut environment of coleopteran larvae. These characteristics, coupled with the broad spectrum of pesticidal proteins identified in the genome, reinforce the potential of INTA Mo4-4 as an effective agent not only against *A. diaperinus* but also against other coleopteran pests [14].

The high-quality genome assembly of INTA Mo4-4 provides substantial insights into its taxonomic placement and functional capabilities. Phylogenomic analyses revealed a close relationship with *B. cereus* and *B. thuringiensis*, with significant genomic similarity to *B. thuringiensis* svar. *morrisoni* BGSC 4AA1, a strain renowned for its coleopteran-targeted activity [16]. The MLST classification of INTA Mo4-4 as sequence type 23 further supports its alignment with coleopteran-active strains [17]. Although the TYGS analysis suggested that INTA Mo4-4 may represent a novel species, the production of proteinaceous parasporal crystals combined with its confirmed insecticidal activity supports its classification within the species *Bacillus thuringiensis* [18].

Importantly, the identification of ten pesticidal protein-coding sequences (CDS) linked to coleopteran toxicity, along with other virulence factors including chitinases and chitin-binding proteins, underscores the potential of INTA Mo4-4 as a biocontrol agent against coleopteran pests through its multiple modes of action. The combination of Cry3 proteins and other pesticidal proteins suggests a multifaceted approach to insect pathogenesis, enhancing its efficacy and reducing the potential for resistance development. The absence of genes associated with thuringiensin biosynthesis, a known insecticidal metabolite, indicates that the strain’s activity relies on protein-based mechanisms rather than secondary metabolites [19].

The findings from this study have significant implications for the development of biocontrol products targeting *A. diaperinus*, and likely, against other coleopterans, which is the subject of future research. The robust genomic and phenotypic profiles of INTA Mo4-4 provide a solid foundation for its application in integrated pest management systems. Its high specificity for coleopteran pests, combined with its genomic similarity to well-characterized *B. thuringiensis* strains, supports its safety and environmental compatibility [14,16]. Moreover, the absence of virulence factors associated with mammalian pathogenesis minimizes concerns about off-target effects.

Our findings align with previous studies on coleopteran-active *Bacillus* strains, particularly those producing Cry3 proteins [16]. However, the broad spectrum of pesticidal proteins identified in INTA Mo4-4 extends its potential application beyond coleopteran pests, encompassing hemipteran and lepidopteran targets as well [14]. This broader activity spectrum positions INTA Mo4-4 as a versatile tool for pest management across diverse agricultural settings.

While this study provides a comprehensive characterization of INTA Mo4-4, several avenues warrant further investigation. Functional assays evaluating the toxicity of individual pesticidal proteins against a range of target pests would elucidate their specific roles and synergistic interactions. Additionally, field trials assessing the efficacy and persistence of INTA Mo4-4 in diverse environmental conditions are essential to validate its practical application.

The observed activity of our strain, nearly four times greater than that of the reference strain, is not yet fully understood but could be related to the strain enhanced ability to produce crystals with a higher proportion of active proteins against *A. diaperinus*. Furthermore, some of the secreted proteins may remain attached to the crystals or spores, similar to what has been reported for Vip3 proteins. Further investigation, including crystal analysis using LC-MS/MS, could help elucidate these hypotheses.

The initial *B. thuringiensis* strain demonstrating specific toxicity to coleopterans was first isolated in 1983 from *Tenebrio molitor* (Tenebrionidae) larvae [20]. This work highlights the importance of conducting screenings to search for new strains with greater activity or a broader host spectrum for the biological control of agricultural pests. Such efforts are crucial to maintaining the biocontrol potential of *B. thuringiensis*, which has sometimes been compromised by the emergence of new insect pests or resistance.

## 4. Conclusions

This study establishes INTA Mo4-4 as a promising candidate for biocontrol of coleopteran pests, particularly *A. diaperinus*. Its unique structural and genomic attributes underscore its potential as a safe and effective alternative to chemical pesticides. The findings provide a valuable framework for the development of innovative biocontrol strategies, contributing to sustainable agricultural practices.

## 5. Materials and Methods

### 5.1. Bacterial strains and production of spore-crystal complexes

*Bacillus thuringiensis* INTA Mo4-4 was isolated from a dust sample of stored products collected in Chacabuco, Buenos Aires, Argentina (S 34° 63’ W 60° 46’). Isolation was conducted using PEMBA medium as described by Holbrook et al. [21]. The strain was subsequently preserved in the Bacterial Stock Collection at the Instituto de Microbiología y Zoología Agrícola (IMYZA) of the Instituto Nacional de Tecnología Agropecuaria (INTA). *Bacillus thuringiensis* svar. *morrisoni* strain *tenebrionis* DSM2803 used as reference was obtained from the Entomopathogen Stock Collection at the Centro de Investigación y de Estudios Avanzados del Instituto Politécnico Nacional (CINVESTAV) in Irapuato, Mexico. Both bacterial strains were cultured in 100 ml of BM broth (composition: 2.5 g NaCl, 1 g KH_2_PO_4_, 2.5 g K_2_HPO_4_, 0.25 g MgSO_4_·7H_2_O, 0.1 g MnSO_4_·H_2_O, 5 g glucose, 2.5 g starch, and 4 g yeast extract per liter, with pH adjusted to 7.2). The cultures were incubated at 30 °C with shaking at 340 rpm for approximately 72 hours, until complete autolysis of the bacterial cells was confirmed under a phase-contrast microscope. For the preparation of spore-crystal complexes, bacterial cultures were centrifuged at 12,000 × *g* for 15 minutes at 4 °C. The resulting pellets were freeze-dried, and the spore-crystal mixture was stored at −20 °C for later use.

### 5.2. Characterization of crystals and their protein composition

Crystals and spores of INTA Mo4-4 were examined using scanning electron microscopy, following a protocol previously described by Benintende et al. [22]. To analyze the protein composition of the crystals, sodium dodecyl sulfate-polyacrylamide gel electrophoresis (SDS-PAGE) was performed [23]. This was done using a 3% stacking gel and a 10% resolving gel in a Bio-Rad Mini Protean 3 Cell system. The electrophoresis was conducted at 50 V for the first 15 minutes, followed by 100 V for 2 hours. Coomassie Brilliant Blue was used for staining, and a high molecular weight standard (Promega) was applied to estimate the molecular weights of the proteins.

To assess the solubilization of spore-crystal mixtures, three different buffers were evaluated: 50 mM Na_2_CO_3_ (pH 10.6), 3.3 M NaBr (pH 6.6), and 0.1 M NaOH (pH 12.9). The spore-crystal suspensions were incubated with the buffers (6:10 vol/vol ratio) overnight at 37°C with gentle shaking. Afterward, the samples were centrifuged at 16,000 × *g* for 10 minutes, and the solubilization efficiency was evaluated by comparing the SDS-PAGE profiles of the supernatants.

For further protein characterization, solubilized crystal proteins from INTA Mo4-4 were treated with trypsin at 37°C, using a 10:1 protein-to-trypsin ratio, with 1 mg of trypsin (EC 3.4.21.4, type IX from bovine pancreas) per ml of deionized water. Samples were incubated for 2, 4, and 6 hours, and the digestion progress was monitored by SDS-PAGE analysis.

### 5.3. Genomic DNA extraction, sequencing, and annotation

Genomic DNA, encompassing both chromosomal and plasmid components, was extracted using the Wizard Genomic DNA Purification Kit (Promega, Madison, WI, USA) according to the manufacturer’s instructions. The purified DNA was visualized via electrophoresis, stained with SYBR Safe dye, and its concentration was determined using a Multiskan SkyHigh µDrop Plate spectrophotometer (Thermo Fisher Scientific, Waltham, MA, USA). Subsequently, an Illumina library was prepared with the extracted DNA, and sequencing was performed in paired-end mode (2 × 150 bp) using Illumina technology at the Unidad Operativa Centro Nacional de Genómica y Bioinformática (ANLIS Malbrán, Buenos Aires, Argentina).

The raw reads were subjected to quality assessment using FastQC v0.12.1 [24]. The genome assembly was performed *de novo* using SPAdes v3.15.4 [25], and the resulting assembly was evaluated for quality using CheckM v1.0.18 [26] and QUAST v4.4 [27] via the KBase platform [28]. The completeness of the genome was assessed using BUSCO v5.5.0 [29], with reference to the “bacilli_odb10” and “Bacillales_odb10” databases.

Species identification and delineation were conducted using the Type Strain Genome Server (TYGS) [30], which calculated digital DNA-DNA hybridization (dDDH) values to determine the closest genomic matches. Additionally, the average nucleotide identity (ANI) between the genome of INTA Mo4-4 and related genomes was computed using the JSpeciesWS online platform [13]. In silico MLST was conducted through the PubMLST website [17]. Genome annotation was performed using the NCBI Prokaryotic Genome Annotation Pipeline [2025 release] and the Rapid Annotation using Subsystem Technology toolkit [RASTtk-v1.073] [31]. Post-annotation, a custom BLAST analysis was performed to identify pesticidal protein genes, employing a curated non-redundant database of insecticidal proteins. Predictions regarding the target insect species susceptible to the identified virulence factors were obtained from the Bacterial Pesticidal Protein Resource Center (BPPRC) specificity database [14].

### 5.4. Insect rearing and bioassay procedures

Specimens of *A. diaperinus* were obtained from the National University of Luján (Argentina) to establish a colony under controlled insectary conditions. The insects were reared in plastic containers measuring 35 x 25 x 10 cm, filled with a mixture of autoclaved commercial chicken feed (Ganave, Argentina) and wheat bran (95:5 ratio). Two slices of fresh carrot, coated with wheat germ oil (Dasipa, Argentina), were added to ensure proper nutrition. The culture was maintained in a growth chamber set at 28 ± 1°C, with a relative humidity of 60-70% and a 14:10 light-dark cycle.

For oviposition, two pieces of corrugated cardboard with grooves were provided as egg-laying substrates, as described by Sallet et al. [11]. These were removed every 48 hours, washed with a 0.3% benzalkonium chloride solution, and placed in separate containers.

Toxicity assays were conducted using the diet incorporation method. Untreated and treated spore-crystal suspensions of INTA Mo4-4 (treated with Na_2_CO_3_ 50 mM, pH 10.6, and solubilized and trypsinized as described in Section 5.2), as well as untreated *B. thuringiensis morrisoni tenebrionis* DSM2803 (6 ml), were added to 34 ml of pre-warmed artificial diet (60°C). The diet consisted of 133.3 g of commercial chicken feed (Ganave, Argentina), 10 g of agar, 2.5 g of ascorbic acid, 1.25 g of sorbic acid, and 2.08 g of nipagin, all dissolved in 1 liter of distilled water. Four hundred microliters of this mixture were dispensed into each well of a 24-well plate (Nunc 143982). A range of spore-crystal concentrations (35-320 µg/ml, dilution factor= 0.65) was also evaluated, with sterile distilled water serving as a control. Mortality of 48 second-instar larvae was assessed for each concentration after a 15-day incubation period at 29°C. Larvae were considered dead if they failed to respond to gentle prodding. The lethal concentration for 50% mortality (LC_50_) was calculated based on data from three replicates using Probit analysis in IBM SPSS Statistics (Version 19).

In addition, INTA Mo4-4 was also evaluated for its ability to produce type I β-exotoxin by counting the number of adult *Musca domestica* that emerged, following the protocol outlined by Sauka et al. [19]. Strains *tenebrionis* DSM2803 and HD-2 were used as a means of comparison and as positive control, respectively.

## Data availability statement

The raw genome sequencing data were submitted to the NCBI with BioSample number SAMN10367088, under BioProject PRJNA503627. The assembled genome is available in the NCBI WGS project, under RIBV00000000.

## Acknowledgments

Leopoldo Palma gratefully acknowledges the Spanish Ministry of Science, Innovation, and Universities, the Spanish State Research Agency, and the European Union for funding his Ramón y Cajal contract (grant ref. RYC2023-043507-I).

