## Supplementary Data S2 for "Unlocking the *Alphitobius diaperinus* (Coleoptera: Tenebrionidae)-combatting capabilities of *Bacillus thuringiensis* INTA Mo4-4 through genomic and phenotypic characterization": Supplementary Data S2_.docx

Biosynthetic gene clusters predicted in the *Bacillus thuringiensis* INTA Mo4-4 genome assembly using the antiSMASH v6.1.1 tool.

| Contig | Type/activity | Location (bp^a^) | Most similar cluster | Similarity (%) |
| --- | --- | --- | --- | --- |
| 1 | RRE-element containing cluster | 93,855 - 111,835 (17,981) |  |  |
| 2 | NI-siderophore | 78,263 - 109,970 (31,708) | Petrobactin | 100 |
| 8 | Non-ribosomal peptide synthetase | 57,884 - 101,854 (43,971) |  |  |
| 9 | Non-ribosomal peptide metallophores, Non-ribosomal peptide synthetase | 9,537 - 61,285 (51,749) | Bacillibactin, NRP | 85 |
| 15 | Terpene | 110,556 - 132,409 (21,854) | Molybdenum cofactor | 17 |
| 16 | Non-ribosomal peptide synthetase | 67,967 - 125,988 (58,022) |  |  |
| 25 | Beta-lactone containing protease inhibitor | 10,568 - 35,806 (25,239) | Fengycin, NRP | 40 |
| 25 | Other unspecified ribosomally synthesised and post-translationally modified peptide product (RiPP) | 86,790 - 97,119 (10,330) |  |  |
| 25 | Other unspecified ribosomally synthesised and post-translationally modified peptide product (RiPP) | 195,195 - 205,455 (10,261) |  |  |
| 40 | NRPS-like fragment | 75,800 - 119,381 (43,582) |  |  |
| 53 | Linear azol(in)e-containing peptides | 10,223 - 33,729 (23,507) |  |  |
| 78 | Non-ribosomal peptide synthetase | 1 - 2,707 (2,707) |  |  |
| 91 | Other unspecified ribosomally synthesised and post-translationally modified peptide product (RiPP) | 2,850 - 10,990 (8,141) | Toyoncin, RIPP | 18 |

^a^The total length of the regions is provided in brackets following the genomic coordinates relative to each contig.
